# A novel model-free feature selection method with FDR control for omics-wide association analysis

**DOI:** 10.1101/2024.03.01.582911

**Authors:** Zi-tong Lu, Xue-ting Song, Yu-fan Gao, Jian Xiao

**Affiliations:** School of Statistics and Mathematics, Zhongnan University of Economics and Law, 430073, Wuhan, Hubei, China

**Author notes:** Membership list can be found in the Acknowledgments section.

## Abstract

Omics-wide association analysis is a very important tool for medicine and human health study. However, the modern omics data sets collected often exhibit the high-dimensionality, unknown distribution response, unknown distribution features and unknown complex associated relationships between the response and its explanatory features. Reliable association analysis results depend on an accurate modeling for such data sets. Most of the existing association analysis methods rely on the specific model assumptions and lack effective false discovery rate (FDR) control so that they may not work well. To address these limitations, we firstly apply a single index model for omics data. This model is free in performance of allowing the relationships between the response variable and linear combination of covariates can be connected by any unknown monotonic link function, and both the random error and the covariates can follow any unknown distribution. Then based on this model, we combine rank-based approach and symmetrized data aggregation approach to develop a novel and model-free feature selection method for achieving fine-mapping of risk features while controlling the false positive rate of selection. The analysis results of simulated data show our method possesses effective and robust performance for all the scenarios. The proposed method is also used to analyze a real ocean microbiome data and identifies some casual taxa unreported by the existing finds.

## Introduction

Advances in high-throughput omics technologies, such as metagenomics sequencing and DNA methylation or protein microarrays, have revolutionized research in medicine. One major use of such technologies is conducting omics-wide association analysis to identify relevant omics features, such as microbial genes and DNA methylation variants, from a large pool of candidate features by analyzing their association with a phenotype of interest, e.g., patient prognosis or response to medical treatment. The identified features can help understand the disease mechanisms, subject to further validation studies, or be used to form more accurate prediction models for personalized medicine [1]. The increasing availability of massive human genomic data sets makes the dimensionality of omics features is much larger than its sample size, also poses new challenges to statistical analysis [2–4]. However, the modern omics data sets collected often not only own the high-dimensionality property but also exhibit unknown distribution response, unknown distribution features and unknown complex associated relationships between the response and its explanatory features. Reliable association analysis results rely on an accurate high-dimensional modeling for such data sets. Such an involved modelling task can be done by based on high-dimensional single index model (SIM) method. The single index model has been the subject of extensive investigation in both the statistics and biology literatures over the last few decades. It generalizes the linear model to scenarios where the regression function is free in performance of allowing the relationships between the response variable and linear combination of covariates can be connected by any unknown monotonic link function, and both the random error and the covariates can follow any unknown distribution. High-dimensional single index models have also attracted interest with various authors studying variable selection, estimation and inference using penalization schemes [5–14]. However, almost of all these methods do not provide a false discovery rate (FDR) controlled multiple testing procedure for simultaneously testing the significance of model coefficients and can not perform FDR controlled feature selection. The multiple testing work of few methods rely on *p*-values with Benjamini and Hochberg (BH) correction [15], and may fail to control FDR in the presence of complex and strong dependence structure among features. Moreover, high-dimensionality of features makes such approach has lower power due to large-scale adjustment burden. Thence developing an effective FDR controlled feature selection method for high-dimensional single index model becomes very desired.

As we know, in the existing literature, the FDR controlled feature selection methods for high dimensional models can be achieved mainly via the following two approaches.

### 1. Knockoff filter-based approach

Barber and Candes [16] firstly introduced the knockoff filter, a new feature selection procedure controlling the FDR in the statistical linear model whenever there are at least as many observations as variables. This method achieves exact FDR control in finite sample settings no matter the design or covariates, the number of variables in the model, or the amplitudes of the unknown regression coefficients, and does not require any knowledge of the noise level. Following the knockoff filter framework, Candes et al. [17] proposed a model-X knockoff FDR control method. However, this method requires both complete knowledge of the joint distribution of the design matrix and repeated derivation of the conditional distributions. While the development of methods to construct exact or approximate knockoff features for a broader class of distributions is a promising area of active research [18, 19], the robustness of how model-X knockoff to the departure of the joint distribution from multivariate Gaussian is currently unknown.

### 2. Symmetrized data aggregation (SDA) approach

To simultaneously test the significance of the regression coefficients in high-dimensional linear regression model, Du et al. [20] firstly proposed a data splitting-based method “SDA” to select features with FDR control. The key idea of the proposed method is to apply the sample-splitting strategy to construct a series of statistics with marginal symmetry property and then to utilize the symmetry for obtaining an approximation to the number of false discoveries. The SDA approach consists of three procedures.

a. The first procedure splits the sample into two parts, both of which are utilized to construct statistics to assess the evidence of the regression coefficient against the null.
b. The second procedure aggregates the two statistics to form a new ranking statistic fulfilling symmetry about zero properties under the null.
c. The third procedure chooses a threshold along the ranking by exploiting the symmetry about zero property between positive and negative null statistics to control the FDR.

Additionally, FDR controlled feature selection procedure can also be implemented by many other statistical inference methods (eg, regression-based modeling, two-sample testing, and statistical causal mediation analysis) [21–26]. However, directly applying these statistical methods to analyze the omics data is usually underpowered and sometimes can render inappropriate results.

In this paper, we employ a general form single index model (SIM) to model omics data. SIM is model-free in performance of allowing the relationships between the response variable and linear combination of covariates can be connected by any unknown monotonic link function, both the random error and the covariates can follow any unknown distribution. Then based on this model, we further utilize rank-based approach [27] and data splitting-based symmetrized data aggregation (SDA) [20] approach to develop a model-free feature selection method for implementing fine-mapping of omics data while controlling the false positive rate of selection. Especially, the proposed method does not rely on *p*-values.

Finally, we design extensive simulation studies to compare the proposed method with the competing methods. The simulation results indicate that the proposed method can successfully control FDR for all the scenarios and further show that in the presence of the sample size is moderate and the errors come from the heavy-tailed Cauchy distribution, or the associated relationship is nonlinear, or the associated relationship is nonlinear with the heavy-tailed Cauchy distribution errors, the proposed method could significantly outperform the competing methods in power performance. For small sample size scenario, compared to the proposed method, other competing methods may either be underpowered or render inappropriate results by having an inflated FDR than the nominal FDR threshold. These results indicate our method owns effective and robust performance for all the scenarios. The proposed method is also applied to the analysis of a real ocean microbiome data set and identifies some casual taxa unreported by the existing finds.

## Materials and methods

This section firstly reviews rank-based approach single index model (SIM) for omics data and then provides parameter estimation methods for SIM. Finally, a multiple testing procedure with FDR control is given for simultaneously testing the coefficients of high-dimensional single index model.

### Review of based on rank approach single index model

Let *X* = (*X*_*i,j*_)_*n×p*_ denote observed *p* omics features matrix on *n* samples; *Y* = (*Y*_*i*_)_*n×*1_ denote the response vector such as disease status, gene expression and so on. Assume that the means of all the *p* features are zero, Rejchel et al. [27] focused on the model-free single index model 1 without intercept

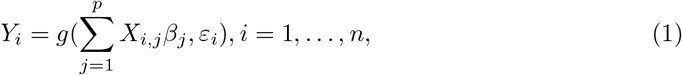

where *β* = (*β*_1_, …, *β*_*p*_)^*T*^ denotes the coefficients associated with the omics features on the response *Y, g*(·) is an unknown monotonic link function. No assumptions are made on the form of the link function *g*, or the distribution of the error *ε*_*i*_, or the distribution of the features *X*_*i,j*_, thence model 1 is model-free. Their purpose is to perform feature selection to identify the feature set *S* = {*j* : *β*_*j*_ *≠* 0, *j* = 1, …, *p* } within the framework of model 1. For this purpose, Rejchel et al. utilized a rank-based lasso approach to sparsely estimate *S* by optimizing the problem

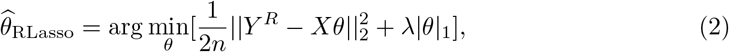

where *η*_*i*_ denotes the random error; 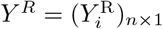 with 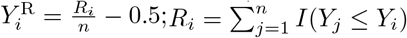 denotes the rank values of the response actual values *Y*_*i*_ by their centered ranks; || · ||_1_ and || · ||_2_ denotes *l*_1_ and *l*_2_ norm, respectively; *λ >* 0 is a tuning parameter of *l*_1_ norm type lasso penalty.

Rejchel et al. [27] highlighted that the rank-based lasso method, as defined in the optimization problem 2, does not estimate true regression coefficients *β* in model 1. However, given some assumptions, they defined a parameter *θ*^0^ by the optimization problem without introducing the penalty

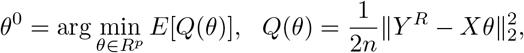

which is related to the true vector of regression coefficients *β*. It is revealed that, under certain standard assumptions, the support 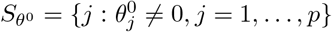 of *θ*^0^ coincides with the support *S* = {*j* : *β*_*j*_ *≠*0, *j* = 1, …, *p*} of *β*. Furthermore, Rejchel et al. [27] demonstrated that the estimator 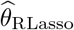 is a consistent estimate of *θ*^0^, and thus can be utilized to identify *S*. More detailed information can be found in the work [27].

### Omics-wide association analysis based on model-free single index model

In real omics data analysis, the response variable *Y* often follows an unknown distribution, which may result in non-linear associated relationships between the response variable *Y* and the linear combination of omics features *X*. In order to simultaneously account for the unknown non-linear associated relationships, unknown distributional response and unknown distributional features, we employ model 1 to model the omics data.

Based on the high-dimensional model 1 with a large number of features (*p* ≥ *n*), we focus on the multiple testing problem 3 under the null hypothesis:

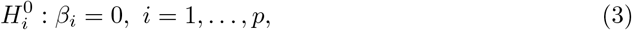

to identify the features associated with the response *Y* while controlling for false positives. Because the above section has shown the support 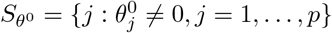 of *θ*^0^ coincides with the support *S* = {*j* : *β*_*j*_ ≠ 0, *j* = 1, …, *p*} of *β* in model 1, the multiple testing problem 3 under the null hypothesis becomes

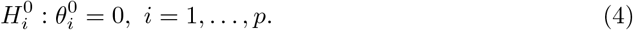

The objective of this paper is to develop a FDR controlled multiple testing procedure for the statistical inference problem 4 to conduct omics-wide association analysis.

### Estimation methods for the parameter *θ*^0^

For the statistical inference problem 4, this section introduces estimation methods for the parameter *θ*^0^ under the high-dimensional model 1 with a large number of features (*p > n*) and the low-dimensional model 1 with a small number of features (*p < n*), respectively. For the high-dimensional scenario, we employ the rank-based lasso method in 2 to obtain the sparse parameter estimates 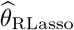. Given the observed variables *X* and the transformed variable *Y* ^*R*^, the optimization problem 2 can be implemented using the R package *glmnet*.

For the low-dimensional scenario, we use rank-based ordinary least squares method to estimate the parameters *θ*^0^ by solving the optimization problem as defined in equation 5

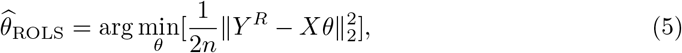

and obtain the parameter estimators

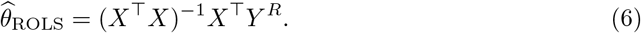

If there are strong correlations among *X*, causing the ROLS method to suffer from multicollinearity and become ineffective, we first utilize the stepwise regression method to eliminate some features. Subsequently, we apply the ROLS method to estimate the parameters of the model constructed by the remaining features. Rejchel et al. [27] demonstrated that the estimator 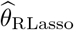 is a consistent estimate of *θ*^0^. From the Theorem 11 [27], it is easy to know 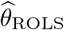 without being thresholded should follow asymptotically normal distribution with the mean being 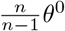 (asymptotical mean being *θ*^0^ as *n* → ∞). More detailed information can be found in the work [27].

### FDR controlled omics feature selection procedure

In this section, we apply the SDA approach [20] illustrated in the “introduction” section to problem 4 for testing the parameter *θ*^0^, and develop a corresponding FDR controlled feature selection procedure. For the two-part independent samples split in the first procedure of SDA, we employ the RLasso and ROLS methods to estimate the parameters, respectively. By the first part sample, we utilize rank-based lasso to sparsely estimate the parameters *θ*^0^ to identify fewer candidate omics features associated with the response. The purpose of reducing feature dimensionality is to alleviate the burden of multiple testing for the third procedure of SDA. Subsequently, for the second part sample, we employ the rank-based ordinary least square method to obtain a more precise estimate under the low-dimensional case. Finally, we utilize the estimates obtained by the two-part samples to construct the FDR-controlled feature selection procedure.

More specifically, the proposed procedure is outlined as follows.

- Step 1: Splitting samples. Given the ratio *γ*, the sample is randomly split into two independent disjoint parts *ζ*_1_ and *ζ*_2_ with sample sizes *n*_1_ and *n*_2_, respectively, where *n*_1_ + *n*_2_ = *n* and *n*_1_*/n* = *γ*. Following the work of [20], we use the ratio *γ* = 2*/*3 to split data for simulation studies and real data analysis.
- Step 2: Selecting the candidate omics feature set using the first part sample *ζ*_1_. The RLasso method is employed to obtain the estimates 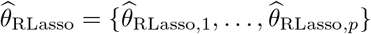 of the parameter *θ*^0^ of *p* features by the first part sample *ζ*_1_. These non-zero estimates are further used to get the candidate feature set 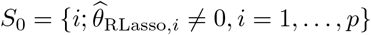. Based on the first part sample *ζ*_1_, we use the candidate features 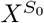 to construct a low-dimensional single index model

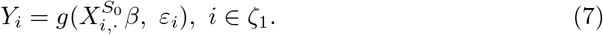 Then the ROLS method is employed for model 7. to obtain the estimates 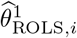 of *θ*_*i*_, *i* ∈ *S*_0_.
- Step 3: Similar to the 2th step, firstly use the candidate features 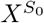, based on the second part sample *ζ*_2_, to construct a low-dimensional single index model, then utilize this model to obtain the estimates 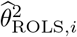 of *θ*_*i*_, *i* ∈ *S*_0_.
- Step 4: Constructing the test statistics under null for problem 4. Under null, we utilize the estimates from the two different part samples to construct the statistics 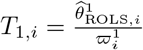 and 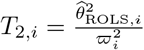, with *i* ∈ *S*_0_ and *T* _2,*i*_ = 0 for *i* ∉ *S*_0_. Here, 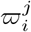 is considered as 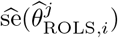 and is viewed as a scaling constant that is independent of 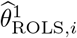 and 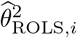. We set the number of re-sampling to be 100 and employ the bootstrap re-sampling method to robustly estimate 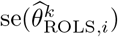 and obtain the estimates 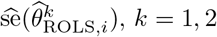.
- Step 5: Aggregating the test statistics.

Aggregate the two statistics obtained in the 4th step to form a new ranking statistic that satisfies the property of being symmetric about zero under the null hypothesis:

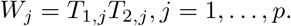

#### Remark 1

Because 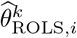 without being thresholded follows asymptotically normal distribution with the mean being 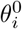, *T*_*k,i*_, *k* = 1, 2 has asymptotically normal distribution with the mean being 0 under null 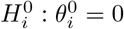. This demonstrates that *W*_*j*_ = *T*_1,*j*_*T*_2,*j*_ possesses the property of being symmetric about zero. Intuitively, the positive and larger *W*_*j*_ values indicate strong evidence against the null hypothesis, while the negative *W*_*j*_ values most likely correspond to null cases.

- Step 6: Choosing the threshold. Given the nominal level *α*∈ [0, 1], a threshold is chosen to exploit the symmetrical property between positive and negative null statistics in order to control the False Discovery Rate (FDR) at the level *α*:

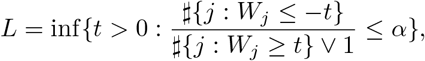

where # denotes the number of elements in a set. The rejected features are given by {*j* : *W*_*j*_ *≥ L*, 1 ≤ *j* ≤ *p*}.
- Step 7: Robustly selecting the discovery set. Suppose we repeat the above 6-step data-splitting procedure *B* times independently. Each time, the set of selected features is denoted as *S*_*b*_, *b* ∈ {1, 2, …, *B*}. For the *j*th feature, we define the empirical inclusion rate 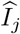 as:

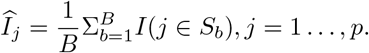

Sort the features based on their empirical inclusion rates in increasing order. Denote the sorted empirical inclusion rates as 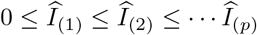. Then select the features 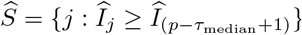, where *τ*_median_ denotes the median size of the selected features over the *B* runs.

#### Remark 2.

The proposed procedure is denoted as SIM-FDR. From the above procedure, FDR control of SIM-FDR method does not rely on *p*-values and requires that the distribution of the aggregated statistics *W*_*j*_ should exhibit symmetry about zero under the null hypothesis. In fact, SIM-FDR does not require that the distribution of *W*_*j*_ must be normal or asymptotically normal. It only requires that the distribution should exhibit the property of symmetry about zero. Thus, SIM-FDR may not heavily rely on the sample size. The simulation results in small sample settings verify this point, indicating that SIM-FDR is more robust than its competitors for a wide range of scenarios, as achieving asymptotic symmetry is much easier in practice compared to achieving asymptotic normality.

## Simulation analysis results

In order to evaluate feature selection performance of the proposed method (SIM-FDR), we consider sample size *n* = 250 and 100 for moderate and small sample size scenarios, and the number of omics features *p* = 400. All simulation settings are replicated 100 times.

### Competing methods

Two methods are considered for comparisons with SIM-FDR.

1. A marginal method that testing one omics feature at a time followed by Benjamini and Hochberg (BH) correction [15], denoting BH method.
2. The original model-X knockoff FDR controlled feature selection method, denoted MXKF method. MXKF method uses based on high-dimensional joint linear regression model approach to analyze continuous response data. It can be implemented by using the R package *knockoffs*.

### Generating omics features

We simulate the *n × p* feature matrix *X* by the multivariate normal distribution *N*_*p*_(*μ*, Σ) with *μ* = (*μ*_1_, …, *μ*_*p*_)^*T*^ and *μ*_*i*_ = 0. The covariance matrix Σ = (Σ_*ij*_)_*p×p*_ is designed as the following structure.

- Σ_*ij*_ = *σ*^|*i* − *j*|^ for *i ≠ j, i, j* = 1, …, *p*, where we set *σ* = 0.5 for moderate strength correlation level among omics features and Σ_*ii*_ = 1 for *i* = 1, …, *p*.

### Simulating the response

We firstly design the regression coefficients *β* = (*β*_1_, …, *β*_*p*_) and then use the coefficients to generate the response. Let the location vector of nonzero values in *β* be

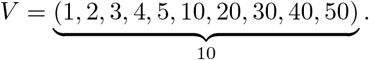

Then we set *β*[*V*], which denotes the nonzero values vector of *β*, to be

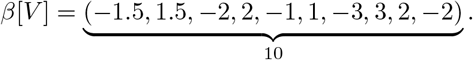

After generating *β*, we employe the following four different type models to simulate the outcome *Y*, where linear regression model is used to simulate linear associated relationships between the response and the omics features, single index model is used to simulate nonlinear associated relationships, and the Cauchy distribution is used to simulate the heavy-tailed distributional random errors.

- Model 1 Linear regression model setting with the normal distributional random error: 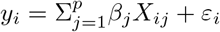, where the error term *ε* is independent of *X* and generated from the normal distribution with location parameter being 0 and the variance parameter being *γ*.
- Model 2 Linear regression model setting with the Cauchy distributional random error: 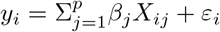, where the error term *ε* is independent of *X* and generated from the Cauchy distribution with location parameter being 0 and the scale parameter being *γ*.
- Model 3 Single index model with the normal distributional random error: 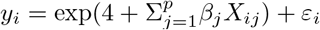, where the error term *ε* is independent of *X* and generated from the standard normal distribution with location parameter being 0 and the scale parameter being *γ*.
- Model 4 Single index model setting with the Cauchy distributional random error: 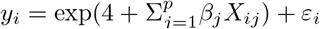, where the error term *ε* is independent of *X* and generated from the Cauchy distribution with location parameter being 0 and the scale parameter being *γ*. In order to consider the different strength association levels between the features and the response, we vary the signal noise ratio (SNR), defined as 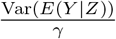 for models 1 and 2, where the scale or variance parameter can be set for models 3 and 4 by the _formula_ 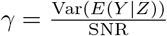.

### Methods settings and comparison measurements

The MXKF method places the burden of knowledge on knowing the complete conditional distribution of *X*, and there is no algorithm that can generate model-X knockoffs for general distributions efficiently [18]. Therefore, we utilize the default design used previously [17] in this simulation. For SIM-FDR methods, the optimal *λ* used in the rank-based lasso are determined through 10-fold cross-validation. For BH method, we test the association between the outcome and each omics feature marginally and apply the BH procedure to these marginal *p*-values to identify significant features. In addition, we set *γ* = 2*/*3 and *K* = 10 for SIM-FDR in simulation analysis.

Given nominal FDR level *α* = 0.05, 0.1, based on 100 simulated data sets, we use empirical FDR and empirical power, defined as

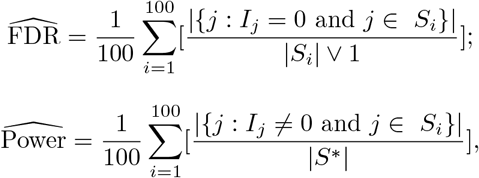

to measure the performance of different methods, where *I*_*j*_ = 0 indicates that the *j*th omics feature is not associated with the response, and *I*_*j*_ ≠ 0 indicates that the *j*th omics feature is truly associated with the response, *S*^*^ = {*j* : *I*_*j*_ ≠ 0, *j* = 1, …, *p*} denotes the indices set of omics features truly associated with the response, |*S*^*^| denotes the number of omics features truly associated with the response, and *S*_*i*_ denotes the indices set of the selected omics feature using the *i*th data set.

### Results for moderate sample size scenario (n=250)

The FDR performance of all the methods is similar for models 1 and 2, and both FDR and power performances of all the methods are nearly the same for models 3 and 4. Consequently, we present the analysis results for models 1 and 2, and models 3 and 4, respectively.

#### Results for models 1 and 2

- FDR performance. For models 1 and 2, as shown in Figs. 1 and 2, the proposed SIM-FDR method effectively controls the actual FDRs at the specified levels for all scenarios. BH method exhibits significantly higher actual FDRs across all scenarios and fails to control the FDR as expected. The MXKF method has significantly higher or lightly higher actual FDRs than the specified FDR levels for all scenarios.

**Fig 1.**
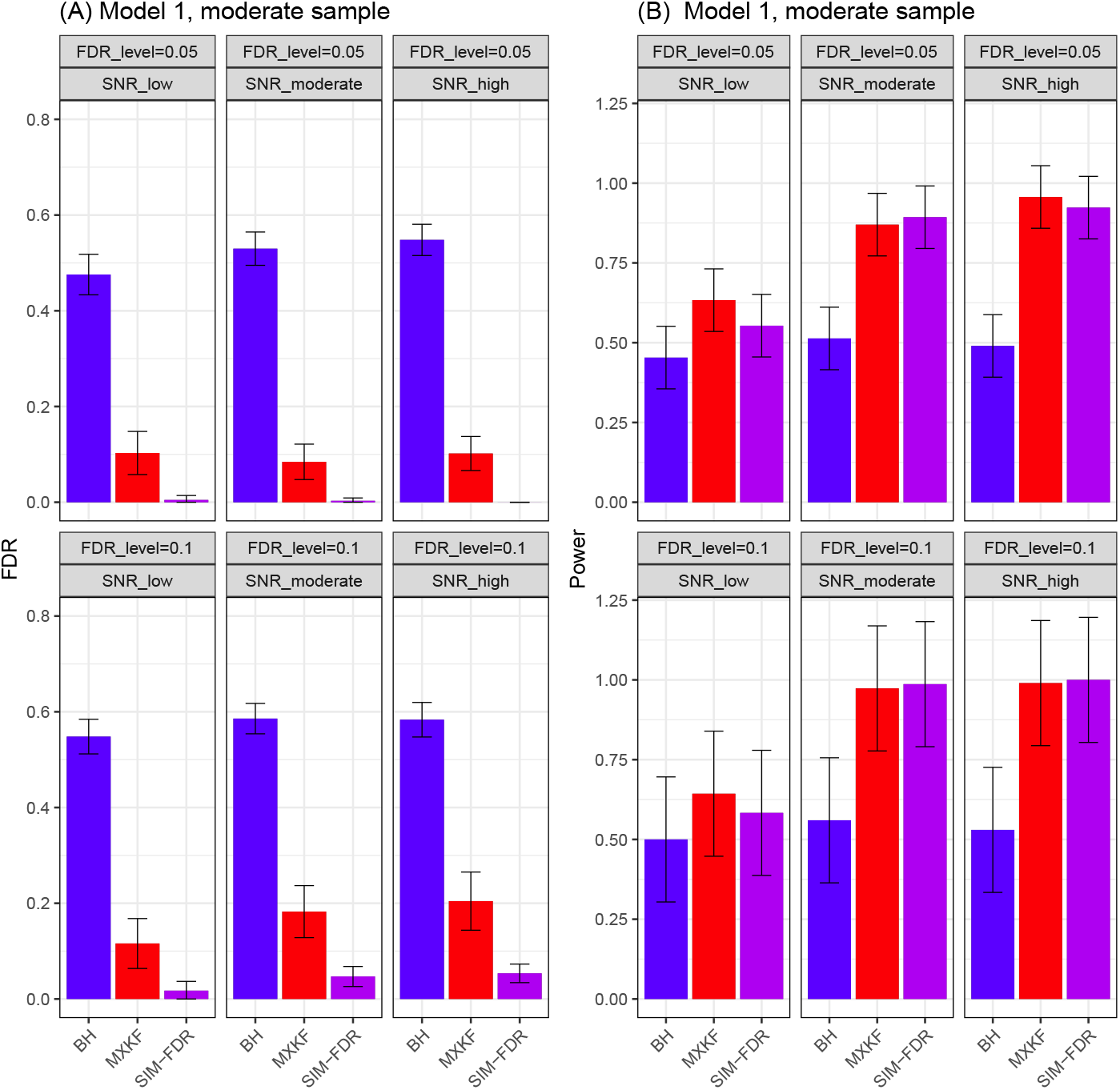
Results for model 1 at moderate sample size case (n=250) FDRs given in left column and powers given in right column are averaged over 100 replications, and their standard deviations (sd) are given on the top of the histogram.

**Fig 2.**
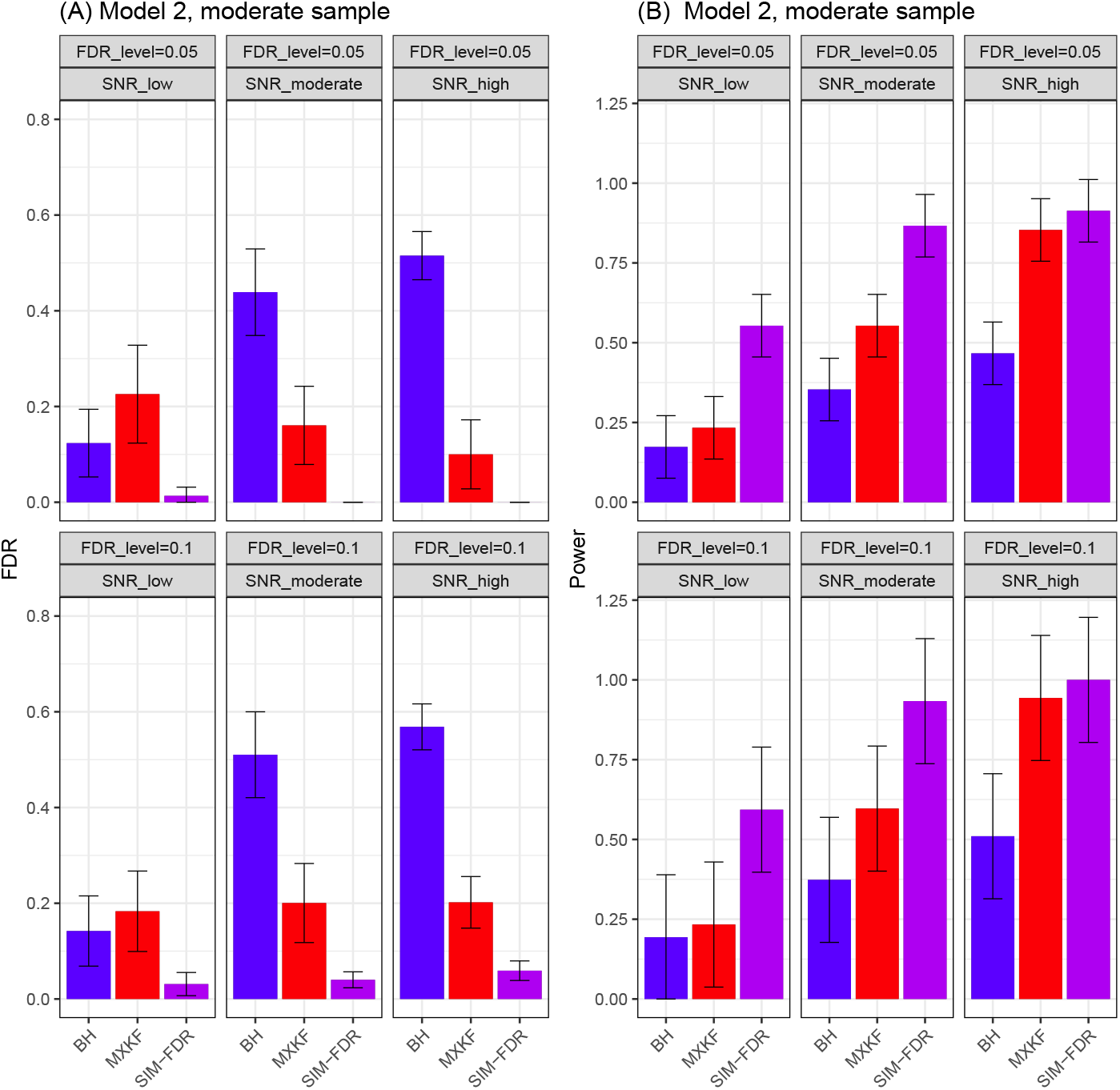
Results for model 2 at moderate sample size case (n=250) FDRs given in left column and powers given in right column are averaged over 100 replications, and their standard deviations (sd) are given on the top of the histogram.
- power performance. For model 1, the MXKF method demonstrates nearly identical power to our SIM-FDR method but at the expense of higher actual FDRs. While for model 2 scenario with the Cauchy distributional errors, SIM-FDR substantially outperforms the MXKF method with the power improvement approaching about 0.36 for some scenarios. BH method possesses too high actual FDRs for these two models so that its performance becomes inconsequential.

#### Results for models 3 and 4

From Figs. 3 and 4, the results for models 3 and 4 demonstrate the same performance for all the methods. Here, we present their results together. For models 3 and 4, SIM-FDR method effectively controls the actual FDRs to the specified levels across all scenarios. In contrast, the MXKF method exhibits much higher actual FDRs for all scenarios and does not achieve the expected level of FDR control. SIM-FDR demonstrates significant performance of MXKF across all scenarios, with power improvement being substantial and approaching about 1.0 for some scenarios.

**Fig 3.**
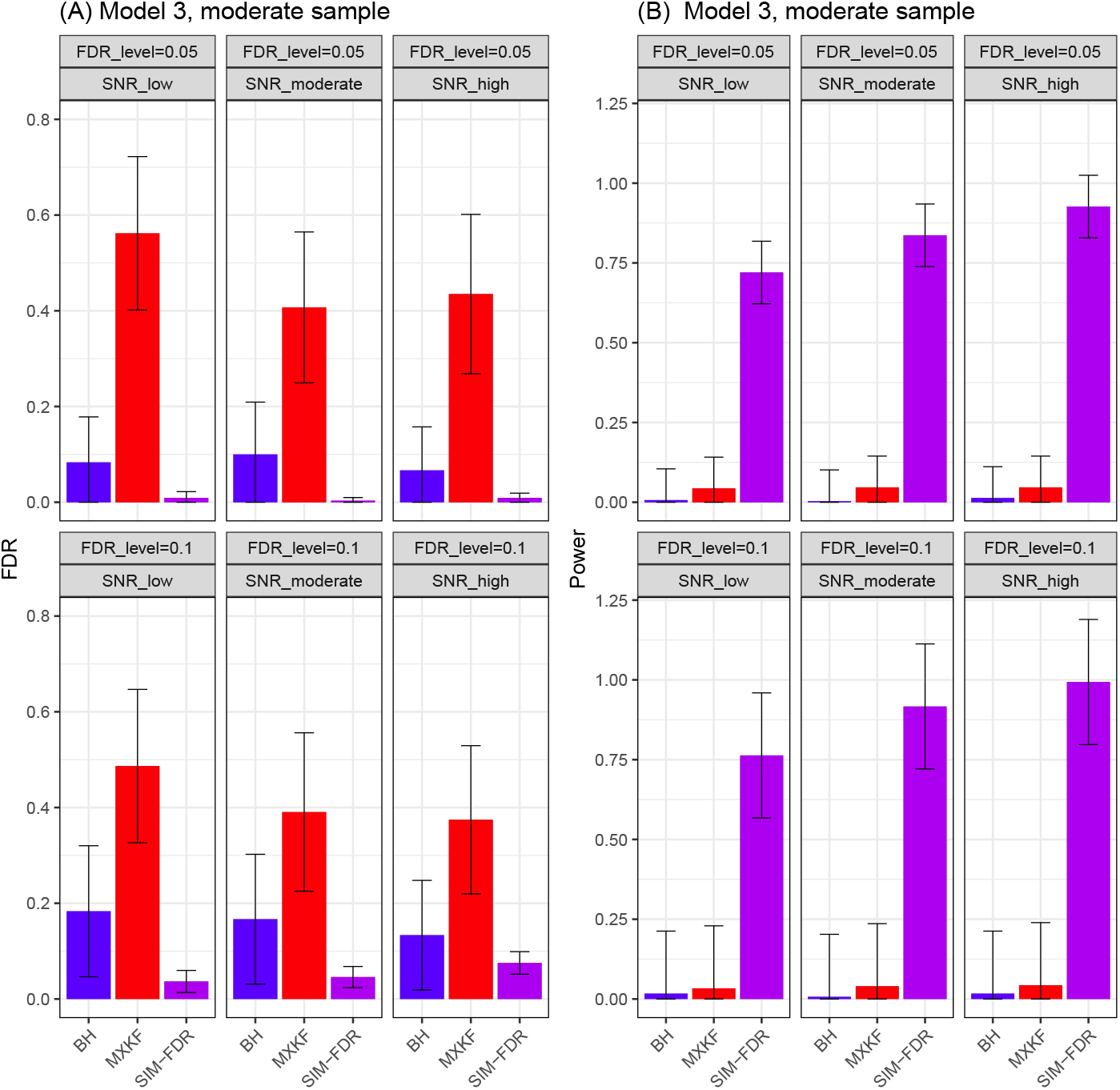
Results for model 3 at moderate sample size case (n=250) FDRs given in left column and powers given in right column are averaged over 100 replications, and their standard deviations (sd) are given on the top of the histogram.

**Fig 4.**
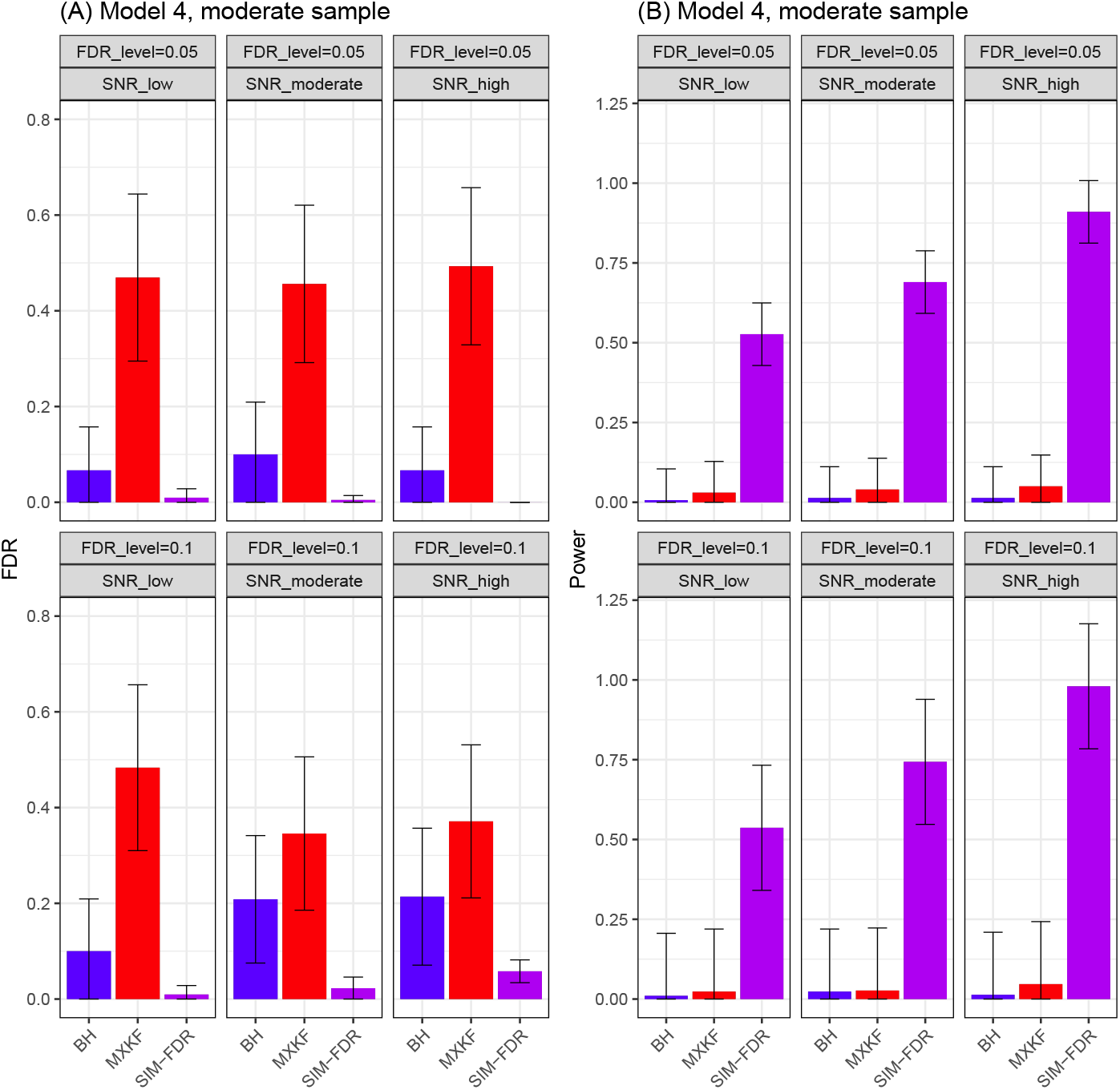
Results for model 4 at moderate sample size case (n=250) FDRs given in left column and powers given in right column are averaged over 100 replications, and their standard deviations (sd) are given on the top of the histogram.

### Results for small sample size scenario (n=100)

The performance of all the methods is similar for models 1 and 2, and models 3 and 4. The analysis results for models 1 and 2, and models 3 and 4 are shown, respectively.

#### Results for models 1 and 2

As shown in Figs. 5 and 6, SIM-FDR method successfully controls the actual FDRs at the specified levels for all scenarios. BH and MXKF methods own significantly higher actual FDRs across all scenarios and can not control the actual FDRs as the desired levels. For the two models, as depicted in Figs. 5 and 6, MXKF method demonstrates slightly higher power than SIM-FDR at the expense of higher actual FDRs.

**Fig 5.**
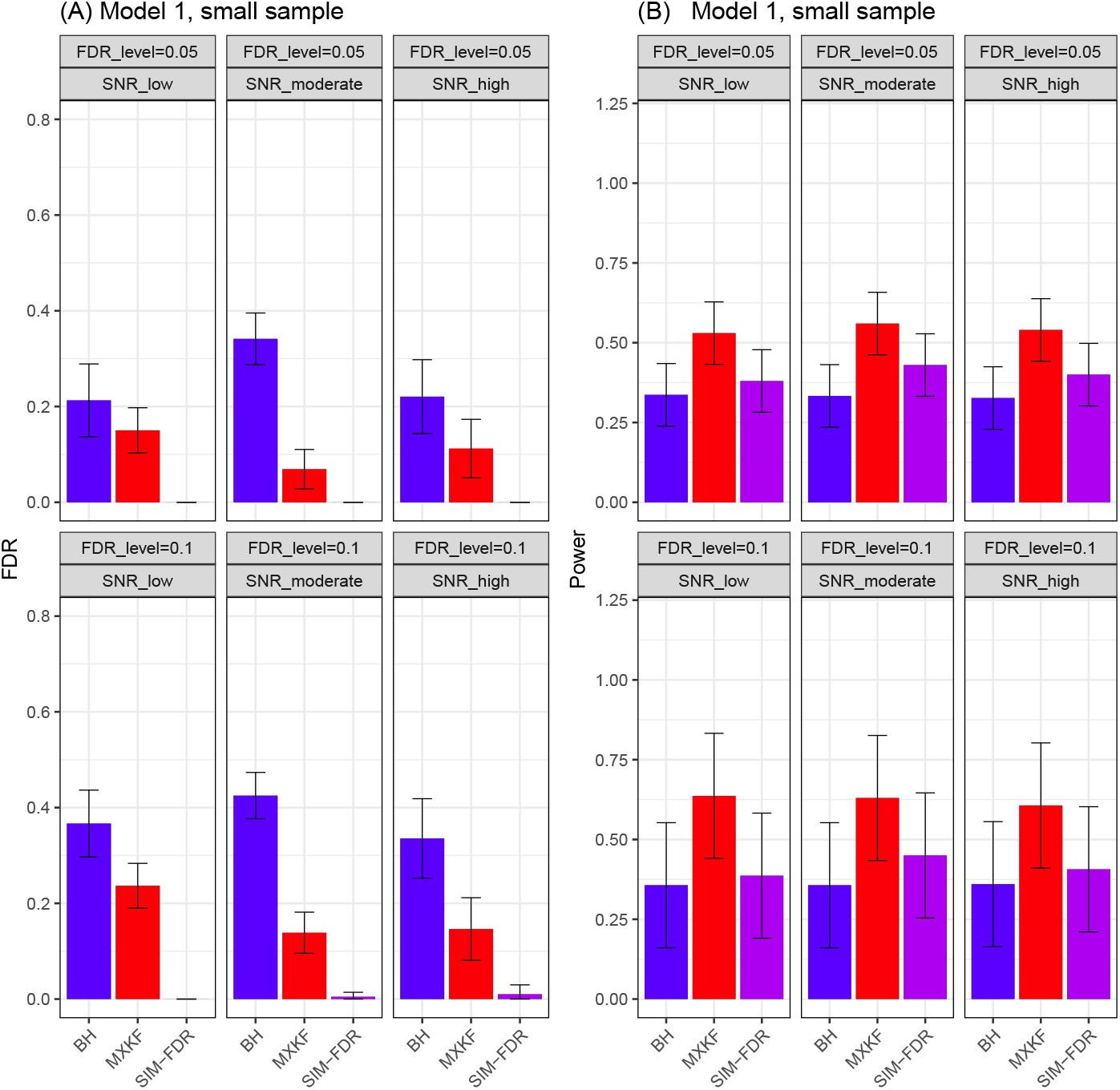
Results for model 1 at small sample size case (n=100) FDRs given in left column and powers given in right column are averaged over 100 replications, and their standard deviations (sd) are given on the top of the histogram.

**Fig 6.**
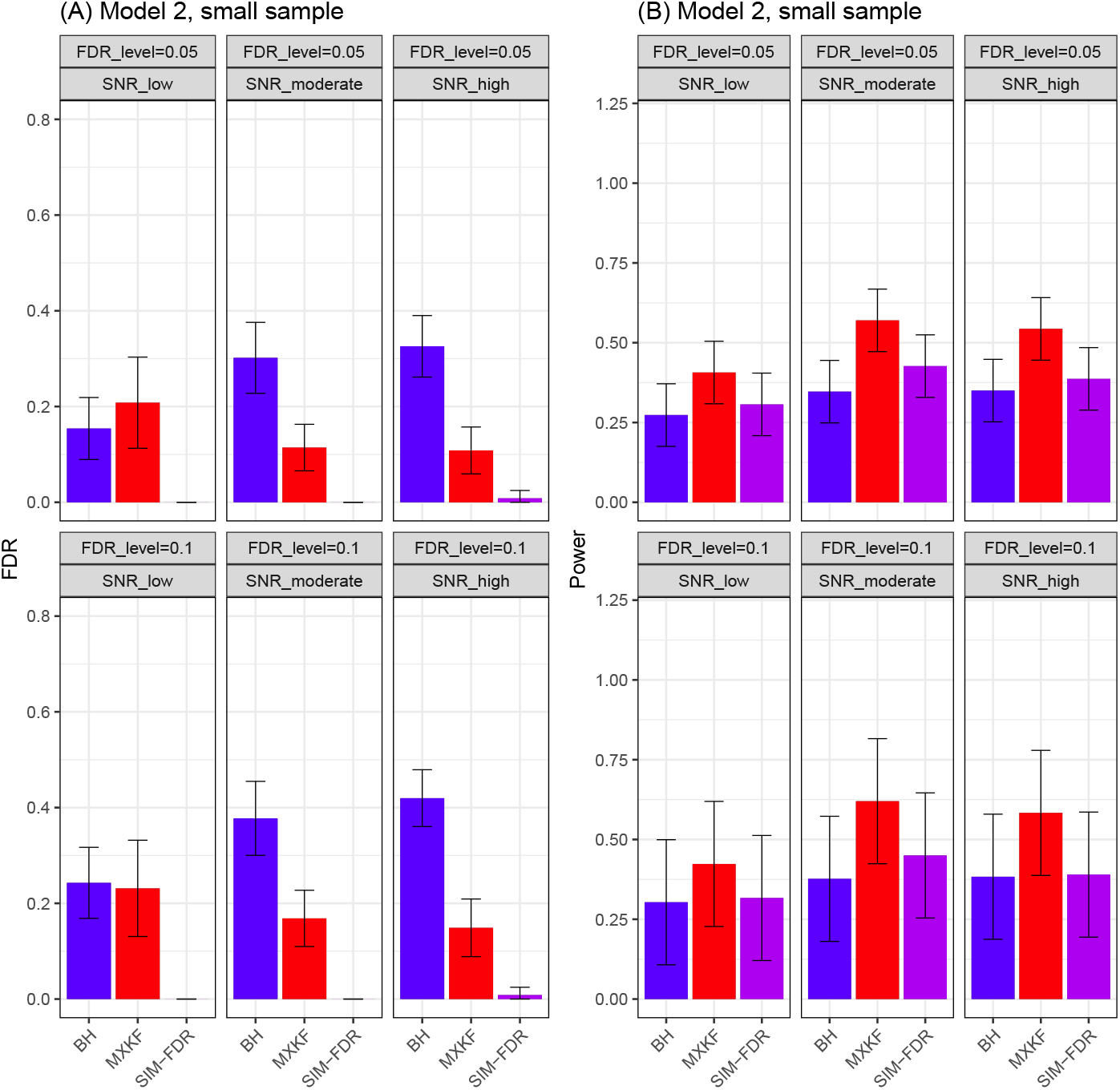
Results for model 2 at small sample size case (n=100) FDRs given in left column and powers given in right column are averaged over 100 replications, and their standard deviations (sd) are given on the top of the histogram.

#### Results for models 3 and 4

From Figs. 7 and 8, the results for models 3 and 4 demonstrate the same performance for all the methods. For models 3 and 4, SIM-FDR method effectively controls the actual FDRs to the specified levels across all scenarios. MXKF method exhibits much higher actual FDRs for all scenarios and does not achieve the expected level of FDR control. SIM-FDR demonstrates significant performance of MXKF across all scenarios, with power improvement being substantial and approaching about 0.38 for some scenarios.

**Fig 7.**
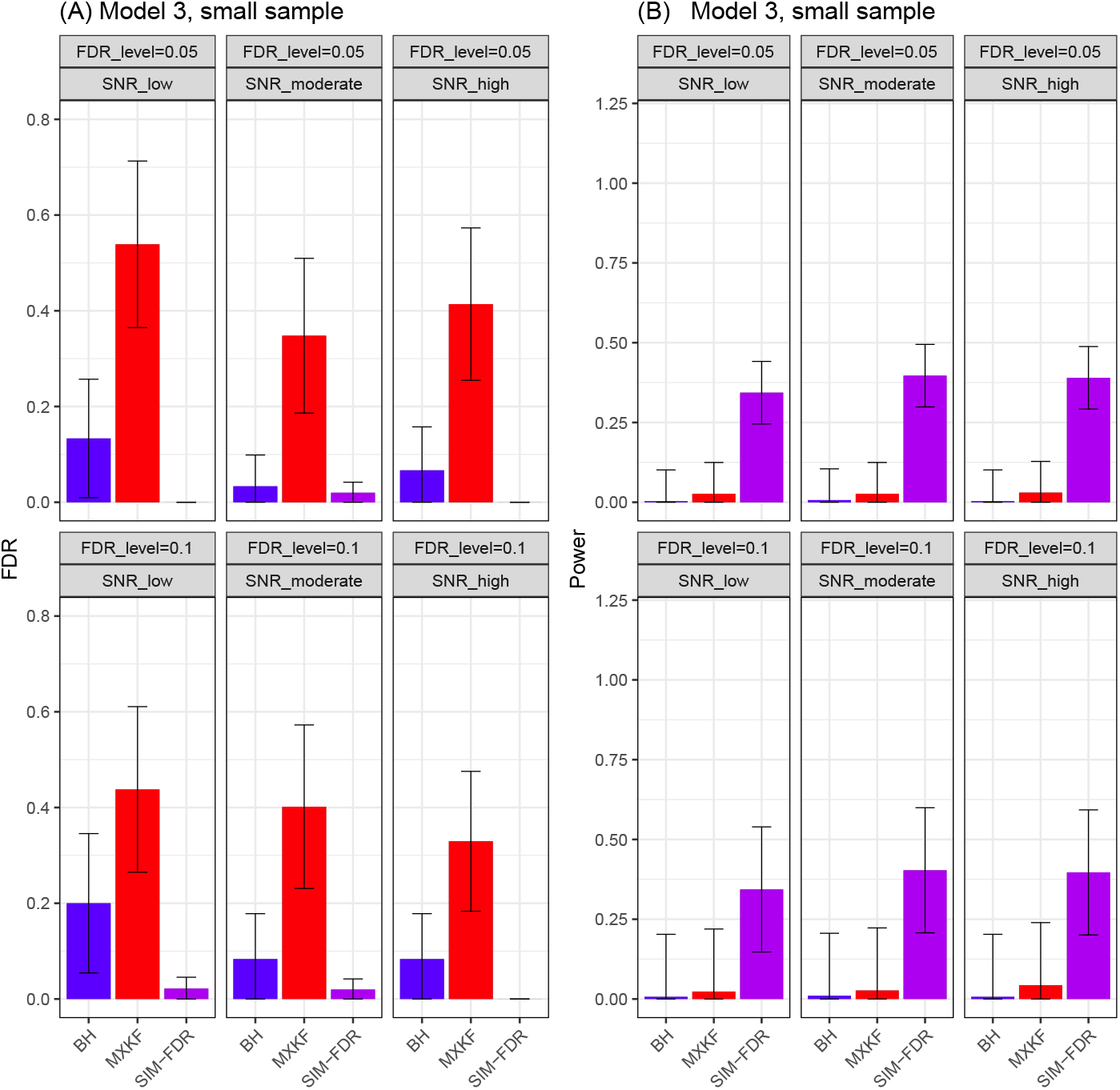
Results for model 3 at small sample size case (n=100) FDRs given in left column and powers given in right column are averaged over 100 replications, and their standard deviations (sd) are given on the top of the histogram.

**Fig 8.**
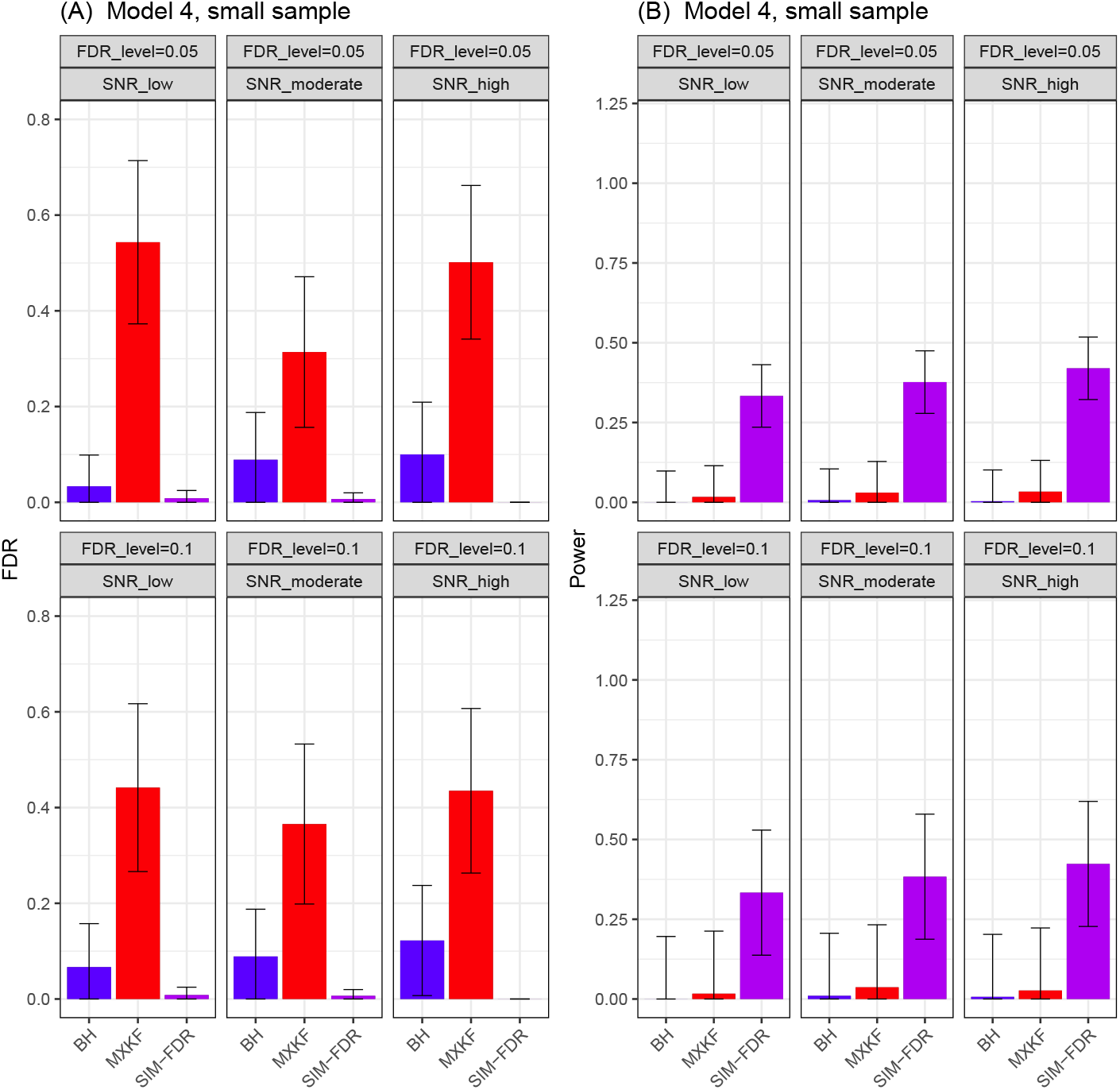
Results for model 4 at small sample size case (n=100) FDRs given in left column and powers given in right column are averaged over 100 replications, and their standard deviations (sd) are given on the top of the histogram.

#### Conclusions of simulating analysis results

Reviewing the results from almost of all the scenarios, the actual FDRs of SIM-FDR are far below the actual FDRs of the other methods. Thence, there should be no need to verify that our procedure has higher power at the price of a higher FDR level. To summarise, compared to SIM-FDR method, other existing methods may either be underpowered or render inappropriate results by having an inflated FDR than the nominal FDR threshold. Such results indicate the proposed SIM-FDR owns robust performance for all the scenarios.

### Real data analysis results

Integrative marine data collection efforts such as Tara Oceans [28] or the Simons CMAP provides the means to investigate ocean ecosystems on a global scale. The collected raw Tara data set is open and can be available at http://ocean-microbiome.embl.de/companion.html. This data set contains *p* = 35651 miTAG OTUs [29] observed on *n* = 139 samples. Using Tara’s environmental and microbial survey of ocean surface water [30], we apply all the methods to identify miTAG OTUs (omics features) associated with some environmental covariates. Especially, salinity is thought to be an important environmental factor in marine microbial ecosystems, then we aimed to identify the miTAG OTUs (omics features) more robustly associated with the response of interest “marine salinity”.

Before applying all the methods, we conducted a series of preprocessing steps to make the Tara data more amenable to the proposed method. Firstly, following the work [31], we calculated read sum of all the 35651 miTAG OTUs (omics features) and removed low-abundance OTUs with read sum less than 10000 reads per sample. We further retained OTUs that appeared in at least 14 samples, resulting in a new OTUs matrix of dimension *n* = 139 and *p* = 1015. Secondly, we normalized OTU raw read counts into composition data with the sum of each row being one. Thirdly, we transformed the composition data using log function and took the log-transformed data as the omics features. Following the analysis of simulated data, we set the same setting *γ* = 2*/*3 for SIM-FDR while let *K* be 30 to stabilize the analysis results of SIM-FDR. We varied nominal FDR levels from 0 to 0.20 for the real data analysis.

The results were presented in Table 1. It is observed that the number of taxa identified by BH method exceeds that of the proposed method SIM-FDR and MXKF, yet MXKF did not identify any taxa fetures for all the FDR levels. This aligns with the simulation results from scenarios: model 1 or model 2, small sample scenario (Figs. 5 and 6), giving the sample size of *n* = 139 and the number of genes *p* = 1015. The Figs. 5 and 6 show that the BH method exhibits higher actual FDRs and fails to control FDR at the given nominal levels. Therefore, the results in Table 1 suggest that BH may yield more false discoveries, while SIM-FDR provides fewer but more precise taxa selection results. Given nominal FDR level 0.20, SIM-FDR method identified six taxa associated with the ocean salinity gradients. From Tables 2 and 3, the identified OTU197, OTU741 and OTU2043 by SIM-FDR come from the class *Alphaproteobacteria*, OTU1473 comes from the class *Deltaproteobacteria*, OTU1439 and OTU520 come from the class *Gammaproteobacteria*. For the Tara data, Bien et al. [31] proposed a tree-aggregated predictive model and also used their method to conduct taxa selection. However, their selection result is very different from that of SIM-FDR. Thus, our results could offer some new perspective of the Tara data.

**Table 1.** The number of selected taxa by all the methods under different nominal FDR levels.

| FDR level | 0 | 0.02 | 0.04 | 0.07 | 0.09 | 0.11 | 0.13 | 0.16 | 0.18 | 0.20 |
| --- | --- | --- | --- | --- | --- | --- | --- | --- | --- | --- |
| BH | 0 | 371 | 444 | 507 | 558 | 588 | 620 | 648 | 671 | 687 |
| MXKF | 0 | 0 | 0 | 0 | 0 | 0 | 0 | 0 | 0 | 0 |
| SIM-FDR | 0 | 4 | 4 | 4 | 4 | 4 | 4 | 5 | 6 | 6 |

**Table 2.** The information of six selected taxa by SIM-FDR at the FDR level 0.20.

| Taxa Name | Rank | Kingdom | Phylum | Class | Order | Family |
| --- | --- | --- | --- | --- | --- | --- |
| OTU197 | Life | Bacteria | Proteobacteria | Alphaproteobacteria | SAR11 clade | Surface |
| OTU741 | Life | Bacteria | Proteobacteria | Alphaproteobacteria | SAR11 clade | Surface |
| OTU2043 | Life | Bacteria | Proteobacteria | Alphaproteobacteria | Rickettsiales | S25-593 |
| OTU1473 | Life | Bacteria | Proteobacteria | Deltaproteobacteria | Desulfuromonadales | GR-WP33-58 |
| OTU1439 | Life | Bacteria | Proteobacteria | Gammaproteobacteria | KI89A clade | f__134 |
| OTU520 | Life | Bacteria | Proteobacteria | Gammaproteobacteria | E01-9C-26 marine group | f__69 |

**Table 3.** The information of six selected taxa by SIM-FDR at the FDR level 0.20.

| Taxa Name | Genus | Species |
| --- | --- | --- |
| OTU197 | g__118 | AY664083.1.1206 |
| OTU741 | g__409 | EU801445.1.1438 |
| OTU2043 | g__681 | JN166192.1.1464 |
| OTU1473 | g__608 | EF574438.1.1503 |
| OTU1439 | g__601 | FR683972.1.1501 |
| OTU520 | g__304 | JF747664.1.1516 |

## Discussion and conclusion

In this paper, we firstly employ a model-free single index model to fit the omics data. Such model is flexible and can account for the nonlinear association relationships with unknown distributional random errors and features. Next, based on this model, we develop an effective false discovery rate (FDR) control procedure for feature selection of high-dimensional single index model, and further use it to fine mapping the omics features while controlling the false discoveries rate of selection. The results on simulated data show when the linear or nonlinear model has heavy-tailed distributional random error at the moderate sample case, the proposed SIM-FDR method could significantly outperform the competing methods in power performance for almost of all the scenarios while controlling FDR very well; while for small sample case, our method can control actual FDRs at the nominal FDR levels for all the scenarios, yet all the other competing methods can not control FDR for almost of all the scenarios. Compared to SIM-FDR method, other methods may either be underpowered or render inappropriate results by having an inflated FDR than the nominal FDR threshold. On the whole, these results indicate SIM-FDR owns robust performance.

However, there has been a wealth of research interest to utilize additional information (eg, phylogenetic information) of microbiome data to increase the power of detection while maintaining FDR control [32, 33]. It is of future interest to incorporate such information in SIM-FDR method framework to further boost the detection power of controlled feature selection.

## Authors’ contributions

J X and ZT L analyzed the data and wrote the paper. YF G prepared the data. XT S reviewed drafts of the paper and wrote the paper.

